# Protein kinase A regulates phosphorylation of UBE2J1 at serine residues S266 in response to glucagon signalling

**DOI:** 10.64898/2026.04.07.716893

**Authors:** Lauryn E. O’Callaghan, Noor D. Algoufi, Dominic S. Dollken, Anwar M. Hashem, John V. Fleming

## Abstract

The ubiquitin conjugating enzyme UBE2J1/Ubc6e localizes to the endoplasmic reticulum where it mediates the ubiquitination and proteasomal degradation of terminally misfolded proteins. Although the protein is known to undergo phosphorylation at serine S184, we have considered modification at an additional site and used a bespoke anti-phospho antibody to confirm phosphorylation also at serine residue S266. Despite the well-described role of UBE2J1 in ER associated degradation (ERAD), we found no evidence for regulation at S266 during Unfolded Protein Response (UPR) induction by thapsigargin. Instead, our studies suggest that phosphorylation occurs independently at the S184 and S266 sites, with mutation at one site failing to disrupt basal phosphorylation at the second. We identified several contexts in which these two phosphorylations were differentially regulated. For example, ER localization, which is important for phosphorylation at S184, was not required for modification at S266, and sensitivity to proteasome inhibitors, which is regarded as a distinguishing feature of the S184 phospho-variant, was unaltered by the S266A mutation. Regarding regulation at S266 on the other hand, we found that pharmacological activation of protein kinase A resulted in rapid phosphorylation, with differential use of phospho-specific antibodies confirming that phosphorylation at S184 was unchanged by this treatment. Hormonal stimulation by glucagon resulted in a similar pattern of UBE2J1 phosphorylation, which occurred exclusively at S266 and could be inhibited by H89. The differential regulation demonstrated in these studies extends our understanding of the UBE2J1 enzyme, and may indicate a role in the integration of energy metabolism with environmental stress conditions.

## INTRODUCTION

UBE2J1 is an E2-ubiquitin-conjugating enzyme anchored to the endoplasmic reticulum via its carboxyl terminus tail. Based on its pattern of cellular localisation and homology with the yeast scUbc6 and human Ube2J2 enzymes it has been attributed a role in ER-associated degradation (ERAD) (1, 2), which is a branch of the Unfolded Protein Response (UPR) that promotes the degradation of misfolded proteins using the ubiquitin-proteasomal pathway (3). As part of ERAD it functions within the translocon complex (4) and, together with the p97/VCP ATP-ase, is involved in the retrotranslocation of misfolded protein substrates through the ER membrane for ubiquitination and degradation by cytosolic proteasomes (3). Interestingly, there is now evidence that UBE2J1 plays an important regulatory role in this process. Not only does it promote degradation of misfolded proteins, but it has also been shown to regulate the steady state expression of several ERAD components (5, 6).

In response to environmental and pharmacological stresses, it is known that UBE2J1 undergoes phosphorylation at serine S184 (7, 8). This can be facilitated by the MK2 kinase that is downstream of p38-MAPK signalling, although in the absence of MK2 (MK2/MK3 deficient MEFS) phosphorylation could be still be detected (7). This suggests that other kinases can regulate this site, and that it represents an integration point for several signalling pathways. In the context of UPR signalling, it has been shown that phosphorylation can occur as a consequence of PERK signalling (although it is not a direct substrate for PERK phosphorylation) (8).

The implications of phosphorylation at S184 remain under active investigation and may be context specific. For example, it does not appear to influence interactions with Parkin, an E3 ligase that partners with it to degrade the Pael receptor (8). In contrast, it increases interactions with the E3 ligase cIAP1 (9) - although the implications of this have not yet been considered in the context of the ubiquitination of misfolded substrate proteins. It is interesting to note that phosphorylation at S184 leads to a less stable form of the enzyme that is targeted for proteasomal degradation (9). Whether cIAP1 is involved in this is not yet known; however, it remains one of the clearest and most detectable functional consequence of phosphorylation at S184.

Although best described for its role in ERAD, several reports have demonstrated the involvement of UBE2J1 in cellular processes beyond the degradation of misfolded proteins (10-16). Considering the intricacies of regulation during ERAD and the expanded range of functional roles in other cellular response processes, it is reasonable to assume that the enzyme is regulate by additional mechanisms. Here we demonstrate that the protein is phosphorylated at serine S266 and explore the impact of this on several parameters of enzyme function and expression.

## MATERIALS AND METHODS

### Plasmid DNA

PCR was used to amplify coding sequences corresponding to full length (amino acids 1 through 318 – 5’) and carboxy-truncated (amino acids 1 through 287) human UBE2J1 were PCR amplified to incorporate a His tag from a human testes cDNA library and cloned into the PstI and Sal I sites of the pEP7-vector as previously described (17), thus generating the pEP7-UBE2J1-His and UBE2J1 ΔTM-His plasmids. The QuickChange protocol (Stratagene) was used to mutate serine residues S184 and S266 of the UBE2J1 coding sequence to alanines, giving rise to pEP7-UBE2J1-S184A-His, UBE2J1-S266A-His and double mutant UBE2J1-S184A/S266A-His plasmids. sHiBiT-GCGR Fusion Vector expressing the human glucagon receptor was a gift from Promega Corporation (Addgene plasmid # 236837 ; http://n2t.net/addgene:236837 ; RRID:Addgene_236837).

### Cell culture and transfections

HEK293T (ATCC) cells were cultured in DMEM supplemented with 10% fetal bovine serum and 100u/ml penicillin/ streptomycin at 37°C in a humidified 5% CO_2_ atmosphere. For transient transfections, cells were seeded a density of 1 × 10^5^ cells per well on 24 well plates pre-coated with poly-L lysine and transfected with Lipofectamine 2000 as described elsewhere (18). Cells were incubated in the presence or absence of pharmacological agents (Mg132, H89 and forskolin) and hormone (glucagon) purchased from Sigma and as described in the figure legends. Cells were harvested using RIPA buffer as described elsewhere (19).

### Immunoblot analysis

Lysates were fractionated on denaturing 11% SDS–polyacrylamide gels for immunoblotting using standard methods. Nitrocellulose membranes were probed with bespoke rabbit anti-UBE2J1, rabbit anti-phospho UBE2J1-pSer184, rabbit anti-phospho UBE2J1-pSer266 (Davids Biotechnologie) or anti-BiP (Cell Signalling Technologies) antibodies. Equal loading of gels was confirmed using mouse anti-α-tubulin or mouse anti-β-actin antibodies (Sigma). All immunoblots shown are representative of at least 3 independent experiments.

## RESULTS

### The UBE2J1 enzymes undergoes phosphorylation at serine residue S266

Previous studies have shown that both endogenous and ectopically expressed UBE2J1 fractionates on SDS-PAGE gels as multiple bands (7, 8). In our transfected HEK293T cell model, we readily detected lower and higher molecular weight bands using an antibody that recognises the total UBE2J1 protein (Fig.1A, middle panel, arrowheads). Development of an anti-phospho antibody that preferentially recognises the S184-phosphorylated protein (anti-pSer184) confirmed that the higher molecular weight band corresponded to phosphorylation at serine S184, and this band was abolished when a phospho-deficient UBE2J1-S184A mutant was expressed (Fig.1A, upper panel).

**Fig. l.**
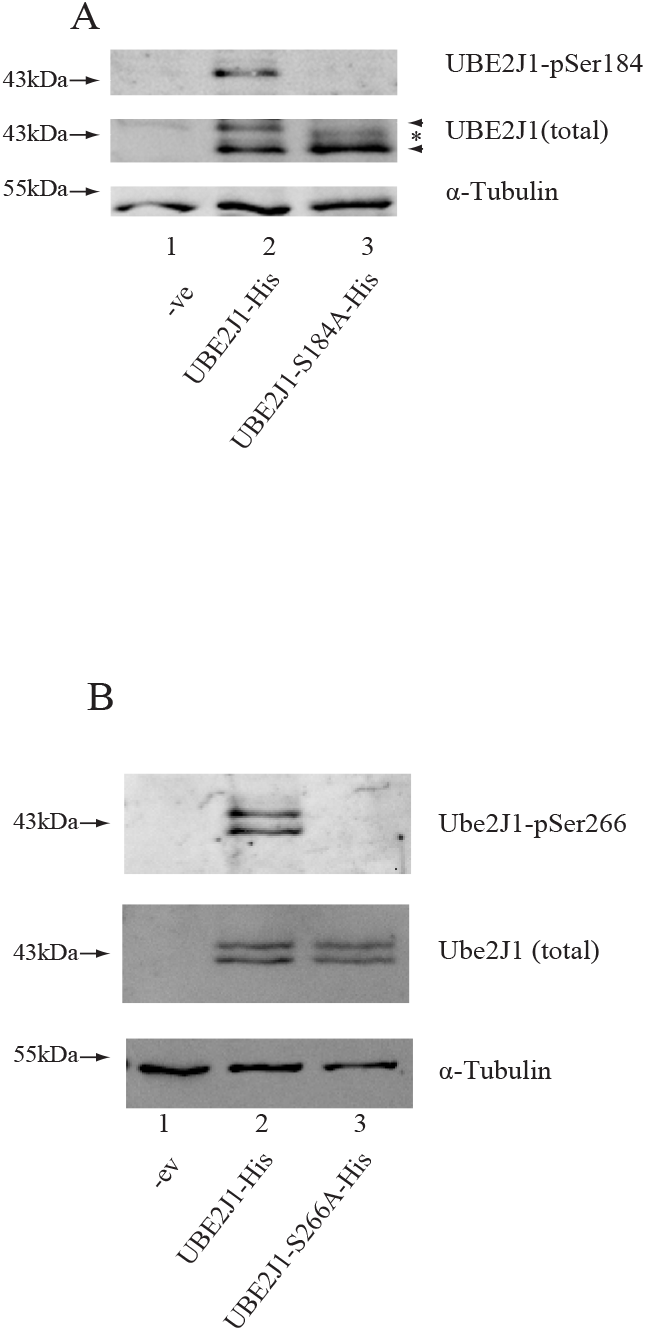
UBE2J1 is phosphorylated at serine residues S184 and S266. (A) HEK293T cells were transiently transfected with plasmids expressing wild type LJBE2J1-His or phospho-deficient UBE2J1-S184A proteins or pcDNA empty vector as a negative control (-ve). Cells were harvested 48 h post-transfection and lysates were analysed on immunoblots using anti-His (UBE2J1-total), or phospho-specific anti UBE2Jl-pSerl84 antibodies. α-Tubulin was used for loading controls. (B) HEK293T cells were transiently transfected with plasmids expressing wild type UBE2J1-His or phospho-deficient UBE2J1-S266A proteins or pcDNA empty vector as a negative control (-ve). Cells were harvested 48 h post-transfection and lysates were analysed on immunoblots using anti-His (UBE2Jl-total), or phospho-specific anti UBE2Jl-pSer266 antibodies. α-Tubulin was used for loading controls. Blots are representative of three independent experiments.

As illustrated in Fig.1A, middle panel, the absence of S184 phosphorylation (ie. the S184A mutant, lane 3) facilitated the observation of an additional weak band (see asterix in Fig.1A), suggesting the possibility of further modifications. Using the NetPhos® kinase prediction software, serine S266 was identified with the highest prediction score (0.998). Mutagenesis of serine 266 to alanine did not result in obvious changes in the pattern of UBE2J1 bands (Fig.1B – middle panel, lane 3). Nevertheless, the development of an anti-phospho S266 antibody (anti-pSer266) demonstrated that the protein does indeed undergoes phosphorylation at this site (Fig.1B – top panel), as specific binding was absent in mutant UBE2J1-S266A lysates in lane 3.

### Phosphorylation at the S184 and S266 sites is mutually exclusive

In lane 2 of Fig.1B, very similar banding patterns were observed when using either the total anti-UBE2J1 (middle panel) or the anti-pSer phospho-specific (top panel) antibodies. At the very least, this observation suggests that both S184-phosphorylated and S184-dephosphorylated proteins can be simultaneously phosphorylated at S266, potentially indicating independent regulatory mechanisms at the two sites. To explore this possibility further, HEK293T cells were transfected to express S184A, S266A or double S184/S266A mutants. As shown in Fig.2, the S266 phosphorylated protein was detectable in the S184A mutant (lane 3). Conversely, mutagenesis of the S266 site to alanine did not prevent the accumulation and detection of the S184 phosphorylated protein (lane 4).

**Fig. 2.**
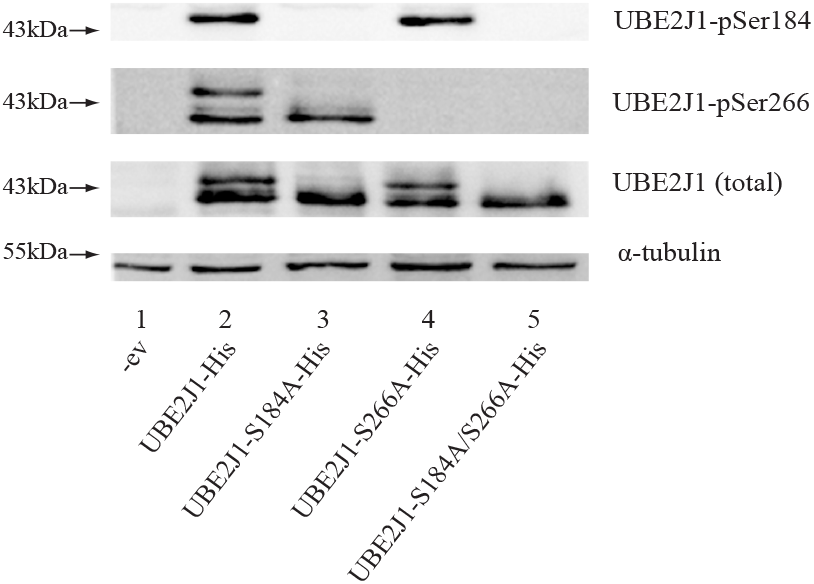
Phosphorylation of VBE2JI at S266 occurs independently of phosphorylation at SI 84 (and *vice* versa) EK293T cells were transiently transfected to express wild type UBE2JI-His, phospho-deficient UBE2J1-SI84A-His, phospho-deficient UBE2J1-S266A-His, or double phospho-deficient UBE2J1-S184A/S266A-His proteins. Cells transfected with pcDNA empty vector were used as negative control (-ve). Cells were harvested 48 h post-transfection and lysates were analysed on inununoblots using anti-His (UBE2JI -total), anti UBE2J1 -pSerl 84 or anti UBE2J1 -pSer266 antibodies as indicated. α-Tubulin was used for loading controls. The blots are representative of 3 independent experiments.

### Phosphorylation of UBE2J1 at S266 does not influence proteasomal degradation of the S184-phosphorylated protein

In terms of enzyme function, increased degradation by the proteasome is recognised as one of the key defining properties of the S184-phosphorylated form of the protein (9). We confirmed these previous reports in our transfected cell model where HEK293T cells expressing UBE2J1 were treated with the proteasome inhibitor Mg132. Our experiments with the anti-phospho pSer184 antibody confirmed the sensitivity of this form of the protein (Fig.3, compare lanes 2 and 3 in the UBE2J1 (total) and UBE2J1-pSer184 panels).

**Fig. 3.**
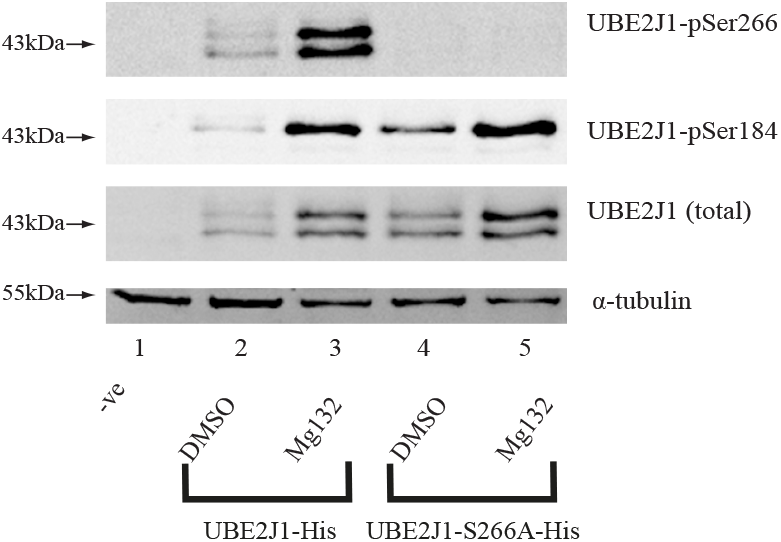
Phosphorylation of UBE2J1 at S266 does not impact on proteasomal degradation of the SI84 phospho-variant. HEK293T cells were transfected to transiently express wild type UBE2J1-His or phospho-deficient UBE2J1-S266A-His proteins. Cells transfected with pcDNA empty vector were used as negative control (-ve). Cells were treated with DMSO vehicle or 20 μM Mg 132 for 16 h before harvesting 48 h post-transfection. Lysates were analysed on immunoblots using anti-His (UBE2J1 -total), anti-UBE2Jl-pSerl 84 or anti-UBE2Jl-pSer266 antibodies as indicated. α-Tubulin was used for loading controls. The blots are representative of 3 independent experiments.

To assess whether S266 phosphorylation is important for this specific property of the S184-phosphorylated protein, western blot membranes were additionally probed with an anti-pSer266 antibody. These studies detected increases in both major UBE2J1 bands (Fig.3, compare lanes 2 and 3 in the UBE2J1-pSer266 panel). While this could be interpreted to mean that S266 phosphorylation contributes to the proteasomal degradation of both S184-phosphorylated and S184-dephosphorylated proteins, the observed response was mirrored by the changes observed for total UBE2J1. Accordingly, it was difficult to assess whether the response was specific to S266 or simply a reflection of the increase in total protein. In parallel cultures where cells were transfected to express the phosphodeficient S266A mutant form of the protein (Fig.3, lanes 4 and 5), it was evident from western blots detecting total and S184-phosphorylated UBE2J1, that the mutation had little or no impact on the Mg132 response.

### Phosphorylation of UBE2J1 at S266 is not dependent on localization to the endoplasmic reticulum or UPR signalling

Previous research has established that the localization of UBE2J1 to the ER is required for S184 phosphorylation (8). The hydrophobic carboxyl-tail region of UBE2J1 (amino acids 288-319) is responsible for this localization pattern. This region facilitates anchoring of the protein into the membrane, positioning the conserved UBC catalytic domain towards the cytoplasm (8, 20). To assess the significance of ER localization for phosphorylation at S266, HEK293T cells were transfected to express a truncated protein lacking the transmembrane domain (UBE2J1-ΔTM). In our transfected HEK293T cell model, we found that the truncated protein retained some limited ability to undergo phosphorylation at S184, consistent with the findings of Oh et al (8). As depicted in the middle panel of Fig.4A(lane 3), the UBE2J1-ΔTM protein continued to produce some of the higher molecular weight bands. Therefore, although we did not observe complete abolition, the vast majority of the protein was present in the S184-dephosphorylated form. This was confirmed when membranes were subsequently probed with the anti-phospho pSerS266 specific antibody (upper panel), revealing that phosphorylation of S266 occurred independently of whether the protein was ER localized or not, with the pattern of expression exactly reflecting the pattern observed for total UBE2J1 protein.

**Fig. 4.**
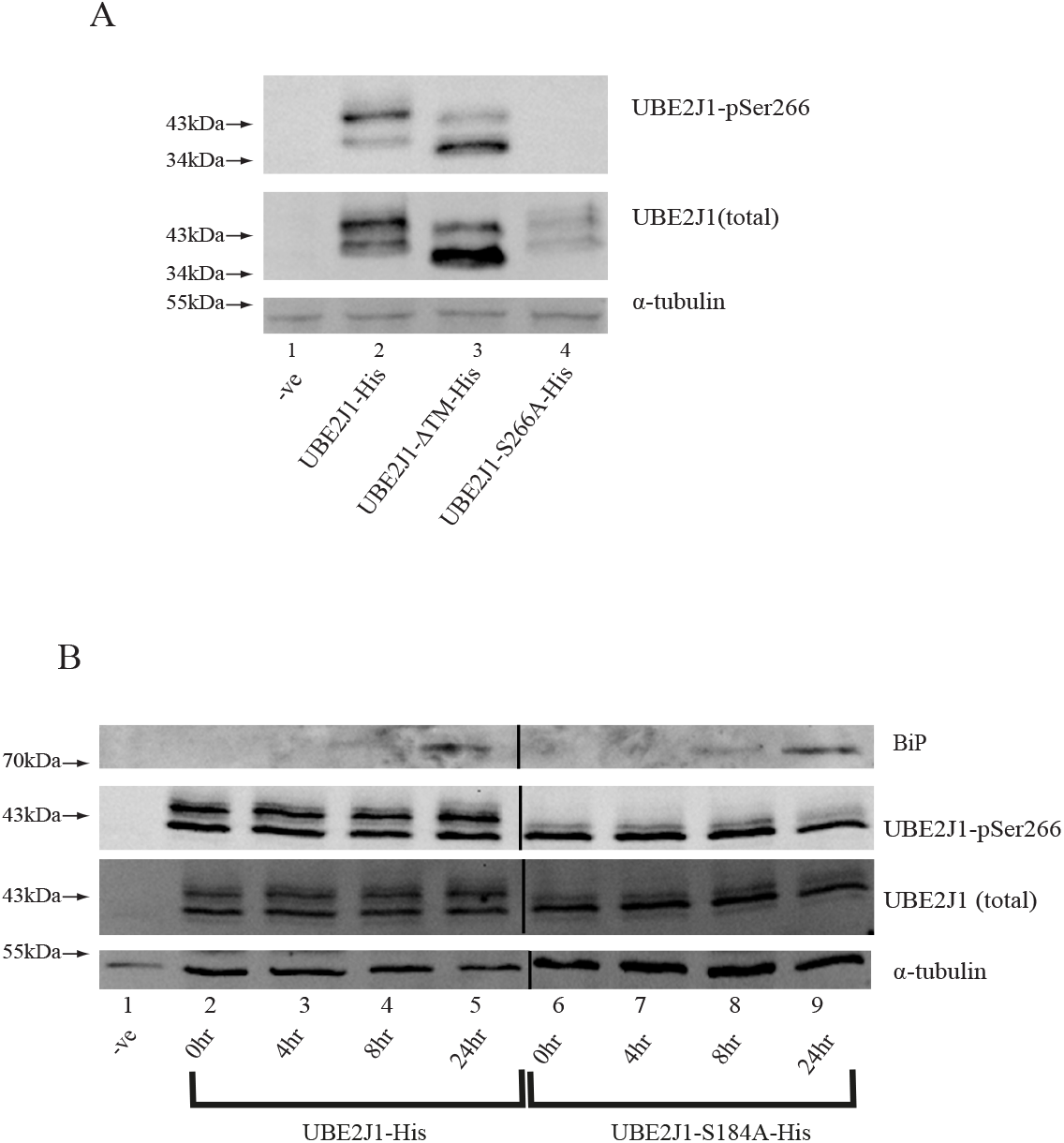
Phosphorylation of UBE2J1 at S266 is unaltered by UPR signalling and does not require ER localization. (A) HEK293T cells were transiently transfected to express wild type UBE2J1-His, truncated UBE2J1-ΔTM-His or phospho-deficient UBE2J1-S266A-His proteins. Cells transfected with pcDNA empty vector were used as negative control (-ve). Cells were harvested 48 h post-transfection and lysates were analysed on immunoblots using an anti-His (UBE2J1-total) or anti UBE2Jl-pSer266 antibodies as indicated, α-tubulin was used for loading controls. The blots are representative of 3 independent experiments. (B) HEK293T cells were transiently transfected to express wild-type UBE2J1-His or phospho-deficient UBE2J1-S184A-His proteins. Cells transfected with pcDNA empty vector were used as negative control (-ve). Cells were treated with 20μM Thapsigargin in reverse time course for the indicated times before harvesting 48 h post-transfection. Lysates were analysed on immunoblots using anti-BiP, anti-His (UBE2J1-total) or anti UBE2Jl-pSer266 antibodies as indicated. α-Tubulin was used for loading controls. The blots are representative of 3 independent experiments. Vertical lines are indicative of where the right-hand side of the blot images was rotated slightly and aligned for presentation purposes.

The UBE2J1 protein plays an important role in ERAD (2, 7). Considering our results in Fig.4A showing that S266 phosphorylation is not tightly linked to ER localization, we examined whether there was any evidence for S266 regulation in response to UPR signalling. HEK293T cells were transiently transfected to express wild type UBE2J1 (Fig.4B, lanes 2-6) or phospho-deficient UBE2J1-S184A (Fig.4B, lanes 7-9), and cells were incubated in the presence or absence of thapsigargin for increasing time periods up to 24 h. Despite increased BiP protein levels that confirmed UPR induction (Fig.4B top panels), we found no evidence of significant regulation of S266 at these time points, and the observed pattern of expression was similar when western blots were probed for either total UBE2J1 protein or phosphorylation at S266 (anti-phospho pSer266).

### Phosphorylation of UBE2J1 at S266 is pharmacologically regulated by forskolin

In contrast, HEK293T cells transfected to express UBE2J1 and stimulated with increasing concentrations of forskolin resulted in the accumulation of the S266-phosphorylated protein. Although the dose-response data shown in Fig.5A were obtained after 3 h of stimulation, subsequent time course experiments using the 10µM dose allowed us to detect an increase in S266 phosphorylation as early as 1 h post-stimulation (Fig.5B). We found no evidence for changes in total UBE2J1 protein levels at this time point, and western blots with the anti-phospho specific pSer184 antibody confirmed that regulation was limited to S266, with no significant changes detected at the S184 site (Fig.5C).

**Fig. 5.**
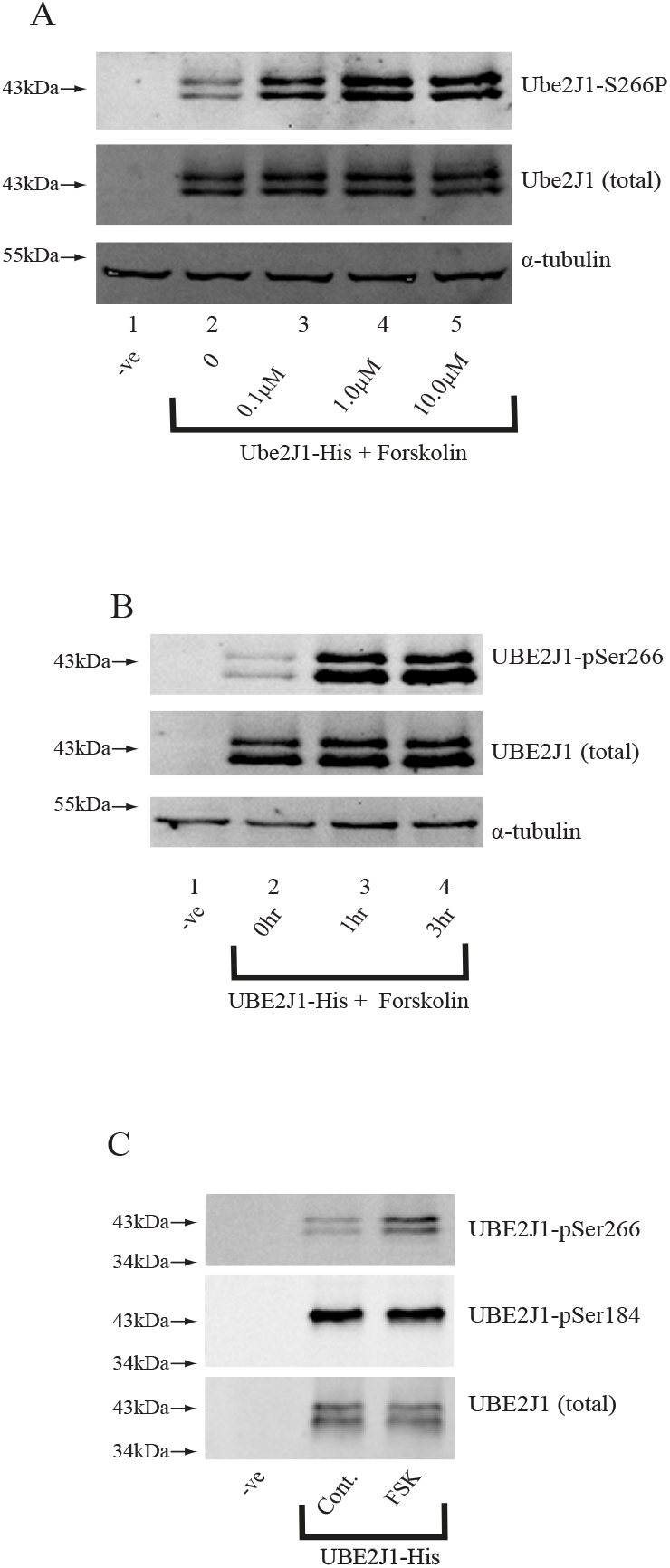
Phosphorylation of IJBE2J1 at S266 is regulated by protein kinase A signalling. (A) HEK293T cells were transiently transfected with a plasmid expressing wild type UBE2J1-His protein or with pcDNA empty vector control (-ve). Cells were incubated for 3 h with DMSO vehicle control (0 - lane 2) or increasing concentration of forskolin as indicated. Cells were harvested 48 h post-transfection and lysates were analysed on immunoblots using anti-His (UBE2J1-total) or anti UBE2Jl-pScr266 antibodies as indicated, a-tubulin was used for loading controls. Blots arc representative of 3 independent experiments. (B) HEK293T cells were transiently transfected with a plasmid expressing wild type UBE2J1-His protein or with pcDNA empty vector control (-ve). Cells were incubated with 10μ M forskolin in reverse time course for the indicated times. Cells were harvested 48 h post-transfection and lysates were analysed on immunoblots using anti-His (UBE2J1 -total) or anti UBE2J1-pSer266 antibodies as indicated, α-tubulin was used for loading controls. Blots arc representative of 3 independent experiments. (C) HEK293T cells were transiently transfected with a plasmid expressing wild type UBE2J1-His protein or with pcDNA empty vector control (-ve). The cells were incubated for 3 h with DMSO (control) or 10μM forskolin (FSK). Cells were harvested 48 h post-transfection and lysates were analysed by immunoblots using anti-UBE2Jl-pSer266 and anti-UBE2Jl-pSerl 84 antibodies as indicated. Anti-His (UBE2J1-total) was used as the loading control. All blots are representative of 3 independent experiments.

Forskolin is a pharmacological regulator of protein kinase A. To further explore the mechanistic basis for the phosphorylation of UBE2J1 at S266, cells transfected to express UBE2J1 were pretreated for 1 h in the presence or absence of the PKA inhibitor H89 before stimulation for 1.5 h in the presence or absence of forskolin. As shown in Fig.6A, treatment of transfected cells with increasing concentrations of H89 resulted in an inhibition of the forskolin induced phosphorylation at S266 (lanes 7-9).

**Fig. 6.**
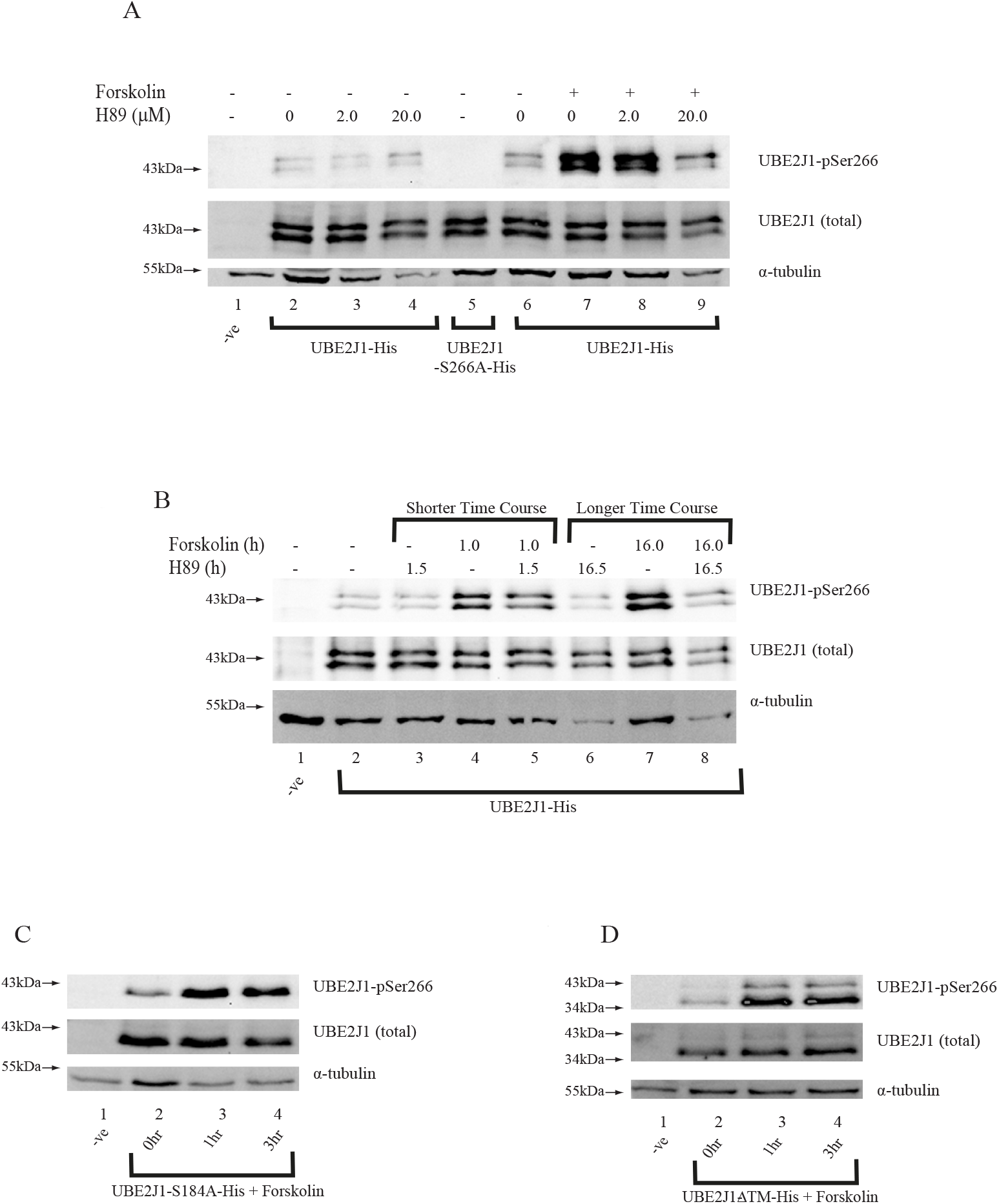
Protein Kinase A stimulated phosphorylation at S266 occurs independently of SI84 and ER localization and is inhibited pharmacologically by H89. (A) HEK293T cells were transiently transfected with plasmid expressing wild type UBE2J1-His protein or with pcDNA empty vector control (-ve). Cells were incubated for 1.5 h in the presence of either DMSO (-) or 10μM Forskolin (+), following a 1.0 h pre-incubation in the presence of either DMSO (-) or increasing concentrations of H89 as indicated. Cells were harvested 48 h post-transfection and lysates were analysed on immunoblots using anti-His (UBE2J1-total) or anti UBE2Jl-pSer266 antibodies as indicated. (B) HEK293T cells were transiently transfected with a plasmid expressing wild type UBE2J1-His protein or with pcDNA empty vector control (-ve). Cells were incubated for short or longer time courses in the presence of DMSO control (-) and cither l0μM Forskolin (+) or 20μM 1189 as indicated. (C and D) HEK293T cells were transiently transfected with plasmid expressing wild type UBE2Jl-S184A-IIis protein (C), UBE2J1-ΔTM-His protein (D) or with pcDNA empty vector control (-ve). Cells were incubated with 10μM forskolin in reverse time course for the indicated times. Cells were harvested 48 h post-transfection and lysates were analysed on immunoblots using anti-His (UBE2JI-total) or anti UBE2Jl-pSer266 antibodies as indicated. α-Tubulin was used for loading control. The blots are representative of 3 independent experiments.

In the absence of forskolin it was noted in these experiments that the phosphorylation of S266 was never completely inhibited by H89 (Fig.6A lanes 2 to 4 compared with lanes 7 to 9). Certainly, it didn’t approach the complete abolition observed for the UBE2J1-S266A mutant (Fig.6A, lane 5). Even when applied over an extended 16.5 h time course (Fig.6B, comparing lanes 6 and 3 with lane 2), the background levels of S266 phosphorylation were retained and remained comparatively unresponsive to H89 treatment. This could reflect differential regulation under basal versus stimulated conditions, with different pathways and kinases responsible. Even if S266 phosphorylation could occur independently of S184 under basal conditions (Fig.4), it remained possible that S184 is important for regulating the S266 phosphorylation response to PKA. To test this, cells were transfected to express the phospho-deficient UBE2J1-S184A mutant and stimulated for increasing time periods with forskolin. As shown in Fig.6C, in the complete absence of S184, we observed an increase in S266 phosphorylation as observed for the wild-type protein. A forskolin induced increase in S266 phosphorylation was also observed when HEK293T cells were transfected to express the truncated UBE2J1-ΔTM protein (Fig.6D). These data confirmed that both basal and PKA regulated phosphorylation at S266 occurs independently of ER localization and S184.

### Phosphorylation of UBE2J1 at S266 occurs in response to glucagon signalling, and is inhibited by PKA antagonists

Considering that S266 phosphorylation was not significantly altered by thapsigargin induced UPR (Fig.4B), we investigated the cellular context it might be regulated. PKA signalling is downstream from many G-protein coupled receptors, including the glucagon receptor that functions alongside the insulin receptor to regulate the energy balance of cells (21). To test whether the phosphorylation of UBE2J1 at S266 constitutes part of the metabolic response, HEK293T cells were co-transfected to express the glucagon receptor, and treated with the hormone for increasing time periods. As can be seen in Fig.7, phosphorylation at S266 was detectable within 30 min, and tended to decrease thereafter through the 2 h timepoint (lanes 2 to 5). This occurred without any regulated change in total UBE2J1 levels and could be inhibited by H89 (compare lanes 5 and 6). The level of phosphorylation was comparable to that observed after 2 h of forskolin stimulation (lane 7). Considering that HEK293T cells are not known to express endogenous receptors, it is noteworthy that the hormonal regulation of UBE2J1 that we observed was dependent on co-transfection with the glucagon receptor, and was not seen in its absence (lane 8).

**Fig. 7.**
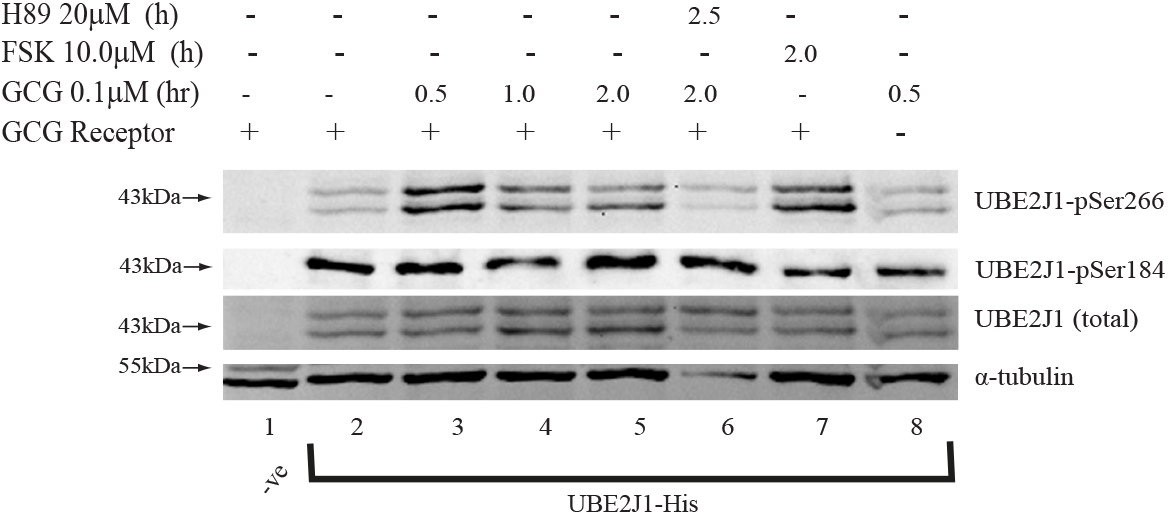
Phosphorylation of UBE2J1 at S266 is regulated by glucagon signalling. HEK293T cells were transiently transfected with a plasmid expressing wild type UBE2J1-His protein or with pcDNA empty vector control (-ve). Plasmid expressing human glucagon receptor (GCG Receptor) or pcDNA empty vector (-) were co-transfected as indicated. Cells were incubated for the indicated time periods in the absence (-) or presence (+) of either 10μM Forskolin (FSK), 0.1μM glucagon (GCG) or 20μM H89. Cells were harvested 48 h post-transfection, and lysates were analysed on immunoblots using anti-His (UBE2Jl-total), anti UBE2Jl-pSerl84 or anti UBE2Jl-pSer266 antibodies as indicated, α-tubulin was used for loading controls. Blots arc representative of 3 independent experiments.

## DISCUSSION

The UBE2J1 enzyme was initially described as a component of the UPR in higher eukaryotes where, as part of ERAD, it functions alongside HRD1 translocon complex components to facilitate the proteasomal degradation of misfolded proteins, such as TCRα (2, 7) and mutant forms of CFTR (22) and α1AT (5). It is not the only ubiquitin conjugating enzyme to play a role in ERAD, but functions instead alongside the Ube2J2 and Ube2G2 enzymes to manage protein misfolding and proteotoxic stress (2, 23). In contrast to these other enzymes, it has been uniquely described as capable of phosphorylation in response to several stresses (7). This raises the possibility that UBE2J1 may play an expanded role compared to other ERAD Ubcs, with opportunities for regulation under different cellular conditions. Here, we used bio-informatic analysis to identify serine S266 as having a high probability for phosphorylation, and employed a transfected HEK293T cell model to describe and characterise phosphorylation at this site. Despite the very clear alterations in electrophoretic mobility that are associated with the mutation of the S184 phospho-site (2, 7-9), we found no such obvious changes when S226 was mutated to alanine. Therefore, the development of the anti-phospho pSer266 antibody was essential to confirm the existence and regulation of the second modified form of the protein.

Using this antibody, we were able to show that phosphorylation at S184 and S266 can occur independently of one another, with mutagenesis of the individual residues failing to abrogate modifications at the second site. It also allowed us to assess whether phosphorylation at S266 could impact S184-associated function. As an example, we explored the accelerated degradation properties associated with S184 modified protein. Although it has been shown that phosphorylation at this site facilitates interactions with the E3 ligase cIAP1 (9), the importance of this on UBE2J1 degradation has not been specifically addressed. Therefore, while it remains unclear why this form is preferentially targeted for degradation, we found that mutation of S266 had little impact, with the wild type and S266A phospho-deficient proteins retaining their sensitivity to proteasome inhibitors.

Our experiments with UBE2J1-ΔTM were also insightful, and although our results confirmed the importance of ER localization for phosphorylation at S184 (we observed decreased accumulation of the S184 phospho-variant consistent with previous reports (8)), we found no evidence to suggest that S266 was being similarly regulated. In the absence of any ER-linked regulation it was not surprising that we failed to observe changes in S266 phosphorylation in response to thapsigargin induced UPR. In this latter case, and despite an increase in BiP levels, there was no obvious change to the S266 phosphorylation status in either the presence (UBE2J1 wild type) or absence of S184 (UBE2J1-S184A).

Considering that S184 phosphorylation may involve multiple kinases and is a feature of multiple stresses (7), it remains possible that simultaneous co-regulation of both S266 and S184 could be a feature under certain cellular conditions. Indeed, we cannot completely exclude co-regulation by other UPR inducers. Certainly, the canonical UPR sensors ATF6, Ire1α and PERK do not appear to be recognised as PKA targets however, there are several points at which cAMP signalling can intersect with the UPR. For example, it is known to influence proteasomal degradation (24, 25), lipid metabolism (26) and vesicular trafficking (27). As well as calcium homeostasis via IP3 receptors (28) and SERCA pumps (29). All of these can impact the UPR and represent opportunities for co-regulation. PKA is also known to regulate immune function, with several points of potential overlap with UPR signalling (30), including regulation by p38 MAPK and LPS (31). Therefore, additional studies are required to assess the possibility of S266 phosphorylation in broader UPR-related functions.

In addition to the cellular response to UPR signalling, recent studies have highlighted that UBE2J1 is not only involved in the management of misfolded and mis-assembled proteins at the ER, but is also involved in several other cellular processes. For example, it functions in the ubiquitination and perinuclear localization of p62 to regulate the expression of the EGF receptor (10), and controls the expression of multiple interferon regulatory factor (IRF) family members during viral infection (11, 14, 15). It has also been implicated in cancer progression through the regulation of RSF3 (13) and the androgen receptor (12), as well as undefined components of the AKT and p53 signalling pathways (16). Therefore, it remains possible that phosphorylation of UBE2J1 at S266 will prove important in regulating these other roles.

In the context of glucagon stimulation and PKA activation there are several scenarios in which protein phosphorylation is coordinated across multiple signalling pathways. For example, glycogen synthase kinase can be regulated by ordered, sequential phosphorylations (32, 33), and for glycogen phosphorylase the first phosphorylation at serine 14 potentiates other modifications (34). Although we found no evidence to suggest that S184 phosphorylation changes in response to glucagon signalling, and that phosphorylation at S266 occurs in the absence of S184 (UBE2J1-S184A), our studies suggest that phosphorylation at S266 might nevertheless be differentially regulated depending on whether it is basal or PKA induced conditions. Even for extended drug treatments, we observed no evidence for complete inhibition by the PKA inhibitor H89. Phosphorylation at S184 is known to exhibit differential sensitivity to pharmacological inhibitors under different cellular conditions. Similarly, we cannot exclude the possibility that S266 phosphorylation acts as a target for several signalling kinases. Certainly, with regard to cellular energy balance, it is noteworthy that UBE2J1 has been shown to play a role in regulating ERAD translocon components that degrade pro-insulin (35). Our studies showing phosphorylation by glucagon may point towards a balancing role, and raise the possibility that UBE2J1 functions also in integrating hormonal regulation of the metabolic state with environmental stress.

## ABBREVIATIONS

Ubc: ubiquitin conjugating
UPR: Unfolded Protein Response
ERAD: endoplasmic reticulum associated degradation
PKA: protein kinase A
GCGR: glucagon receptor

## ACKNOWLEDGEMENTS

The authors wish to thank the Ministry of Education, Saudi Arabia for funding support for Noor Algoufi, and T.M. Wac for useful suggestions.

